# Paternal methamphetamine exposure induces higher sensitivity to methamphetamine in male offspring through driving ADRB1 on CaMKII-positive neurons in mPFC

**DOI:** 10.1101/2022.08.19.504512

**Authors:** Yanyan Zheng, Dekang Liu, Hao Guo, Wenwen Chen, Zhaoyu Liu, Zhaosu Li, Tao Hu, Yuanyuan Zhang, Xiang Li, Ziheng Zhao, Qinglong Cai, Feifei Ge, Yu Fan, Xiaowei Guan

**Author notes:** These authors are equally contribute to this work (Yanyan Zheng, Dekang Liu). Correspondence to: Prof. X Guan, Dr. Y Fan.

## Abstract

Paternal abuse of drugs, such as methamphetamine (METH), elevates the risk of developing addiction in subsequent generations, however, its underlying molecular mechanism remains poorly understood. Male adult mice (F0) were exposed to METH for 30 days, followed by mating with naïve female mice to create the first-generation mice (F1). When growing to adulthood, F1 were subjected to conditioned place preference (CPP) test. Subthreshold dose of METH, insufficient to induce CPP normally, were used in F1 (METH^F1^). Selective antagonist (betaxolol) for β1-adrenergic receptor (ADRB1) or its knocking-down virus were administrated into mPFC to regulate ADRB1 function and expression on CaMKII-positive neurons. METH-sired male F1 acquired METH^F1^-induced CPP, indicating that paternal METH exposure induce higher sensitivity to METH in male F1. Compared with saline (SAL)-sired male F1, CaMKII-positive neuronal activity was normal without METH^F1^, but strongly evoked after METH^F1^ treatment in METH-sired male F1 during adulthood. METH-sired male F1 had higher ADRB1 levels without METH^F1^, which was kept at higher levels after METH^F1^ treatment in mPFC. Either inhibiting ADRB1 function with betaxolol, or knocking-down ADRB1 level on CaMKII-positive neurons (ADRB1^CaMKII^) with virus transfection efficiently suppressed METH^F1^-evoked mPFC acyivation, and ultimately blocked METH^F1^-induced CPP in METH-sired male F1. In the process, the p-ERK1/2 and ΔFosB may be potential subsequent signals of mPFC ADRB1^CaMKII^. The mPFC ADRB1^CaMKII^ mediates paternal METH exposure-induced higher sensitivity to drug addiction in male offspring, raising a promising pharmacological target for predicting or treating transgenerational addiction.

**HIGHLIGHTS:**

- Paternal methamphetamine (METH) exposure induces higher sensitivity to
METH in male F1 during adulthood, accompanied with higher ADRB1 level in mPFC.
- METH use in F1 (METH^F1^) evokes more CaMKII-positive neurons in mPFC of METH-sired than saline-sired male F1.
- Inhibiting ADRB1 function or knocking-down ADRB1 level on CaMKII-positive neurons (ADRB1^CaMKII^) efficiently suppresses METH^F1^-evoked mPFC activation, and ultimately rescues transgenerational susceptibility to addiction in male F1.

## INTRODUCTION

Paternal psychostimulants intake elevates the risk of developing addiction in subsequent generations [1, 2], which remains major health concern. Methamphetamine (METH) is a commonly abused psychostimulant worldwide. One recent study found that METH-sired rats exhibit more sensitized behaviors in response to acute drug use [3], indicating that paternal METH use may enhance susceptibility to drugs in offspring. The medial prefrontal cortex (mPFC) is critically implicated in the processes of METH addiction [4, 5] and susceptibility to drugs [6–8]. Human imaging studies show that prenatal METH exposure significantly affect the volume, cortical thickness and connection of offspring brains, especially the prefrontal cortex (PFC) [7, 9, 10]. Previously, we found that mPFC were blunted to stimulants in METH-sired mice during adulthood [11]. However, the intrinsic molecular mechanism of mPFC underlying transgenerational effects of paternal METH abuse on offspring remains much to be clarified.

Adrenergic innervation on mPFC glutamategic (CaMKII-positive) neurons plays essential roles in neurophysiology and neuropathology. The β1-adrenergic receptor (ADRB1) is highly expressed on mPFC CaMKII-positive neurons (ADRB1^CaMKII^), which has been reported to participate in cognition [12, 13], reward [14], arousal [15], motivation [16], and stress [17]. Generally, activating ADRB1 triggers neuronal excitability within PFC [18]. In animal models of cocaine-induced conditioned place preference (CPP), a model to evaluate reward and cue-related stimulations of drugs, ADRB1 administration efficiently blocks the reinstatement behaviors to cocaine in mice [19]. Prenatal events exposure, malnutrition for example, impairs the long-term potentiation in PFC of offspring, which leads to memory deficits [20–22], but could be fully recovered by ADRB1 activation [23]. Our previous transcriptomic analysis screened out a higher levels of *Adrb1* gene in mPFC of METH-sired male F1 mice (not published). These findings suggest that ADRB1 might involve in drug-related transgenerational neurotoxicity, such as mPFC plasticity and vulnerability of addiction.

The β-adrenergic innervation could regulate subsequent signals though G protein-dependent or/and β-arrestin-dependent pathway. As one of classical G protein coupled receptors (GPCRs), ADRB1 has been well-studied to be coupled to Gs protein which lead to an increase in protein kinase A (PKA)-dependent signaling pathway including cAMP, PKA, and cAMP response element-binding protein (CREB) [12, 24]. Recent reports have revealed that several ligands called “biased ligands”, such as β-adrenergic receptors, elicit G protein-independent and β-arrestin-dependent pathway [25, 26], which may be served as negative feedback of the G protein-dependent pathway [27]. Huang et al. [28] find that β-Arrestin-biased β-adrenergic signaling promotes extinction learning of cocaine reward memory. At present, the price signaling pathway of ADRB1 in drug-preferred behavior remains much to be clarified.

In the present study, male adult mice (F0) were exposed to METH for 30 days, followed by mating with naïve female mice to create the first-generation mice (F1). When growing up to adult age, F1 mice were subjected to conditioned place preference (CPP) test to evaluate sensitivity to METH. Subthreshold dose of METH, that is insufficient to induce CPP normally, were used in F1 mice (METH^F1^). The role of ADRB1^CaMKII^ in transgenerational susceptibility of addiction were explored in METH-sired F1 mice.

## MATERIALS AND METHODS

### Animals

Male C57BL/6 wild type (WT) mice weighing 22-25 g were used. All animals were housed at constant humidity (40~60%) and temperature (24 ± 2°C) with a 12-hour light/dark cycle and allowed free access to food and water. The animals were handled for three days prior to onset of experiments. All procedures were carried out in accordance with the National Institutes of Health Guide for the Care and Use of Laboratory Animals of China and approved by the Institutional Animal Care and Use Committee (IACUC) at Nanjing University of Chinese Medicine.

### Drug treatment and tissue collection

Male C57BL/6 WT mice were assigned to administration of METH (5 mg/kg, dissolved in saline, i.p.) or saline (0.2 mL, i.p.) once daily at 10 a.m. for 30 consecutive days. At 24 h after the last injection, the F0 mice were mated with three naïve female C57BL/6 WT mice for 4 days to create METH-sired or saline-sired first-generation (F1) offspring mice.

On postnatal day 21 (PD21), male and female F1 mice were weaned and kept (4 male mice per cage) in homecages. On PD60, the mPFC tissue were collected from some male F1 mice. From PD70, some male and female F1 mice were subjected to METH-induced CPP training and test. In this study, to examine the sensitivity to METH, subthreshold dose of METH (0.25 mg/kg) were used to induce CPP in male F1 mice. Betaxolol, a specific β1-adrenergic receptor (ADRB1) antagonist (Cat# S2091, Sellekchem, USA) was diluted to 0.6 μg/μl in sterile saline. Brains of male F1 mice were collected on PD60 and PD73. The brains of PD60 mice were collected without any treatment to represent baseline condition. The brains of PD73 mice were collected within 60 min after CPP test.

Some male naïve mice were only subjected to saline or METH treatment. On Day 1 (D1) and D2, mice were injected with saline (0.2 mL, i.p.) in the morning, and saline (0.2 mL, i.p.) or METH (0.25 mg/kg, i.p.) in the afternoon once daily at homecage. Their mPFC tissue were collected on D3 within 60 min of the last injection.

### CPP

In the present study, the 0.25 mg / kg dose of METH administration (METH^F1^), being insufficient to produce CPP in naïve mice (subthreshold dose), was used to induce CPP in male F1 mice and female F1 mice. The CPP test was performed in the TopScan3D CPP apparatus (CleverSys, VA, USA), which is constructed of two distinct chambers (15 × 15 × 23 cm each) separated by a removable guillotine door. The CPP procedure consisted of three phases including the pre-conditioning test (Pre-test, PD70), conditioning (CPP training, PD71-72) and post-conditioning test (CPP test, PD73). During CPP training, both METH-sired and SAL-sired F1 mice were injected with saline (0.2 mL, i.p.) in the morning or METH (0.25 mg/kg, i.p.) in the afternoon once daily. After each injection, the mice were confined to one conditioning chamber (SAL-paired chamber or METH-paired chamber) for 45 min, and then returned to their homecages. During the Pre-test and Test, mice freely access two chambers for 15 min. The CPP score is the time spent in METH-paired chamber minus that in SAL-paired chamber, and the ΔCPP score is the CPP score of Test minus the CPP score of Pre-test.

### Immunofluorescence

Briefly, the brains were perfused with 4% paraformaldehyde (PFA), and coronal brain sections (30 μm) were cut on a cryostat (Leica, Germany). The sections were incubated in 0.3% (v/v) Triton X-100 for 0.5 h, blocked with 5% donkey serum for 1.5 h at room temperature, and incubated overnight at 4°C with the following primary antibodies: rabbit anti-c-Fos (1:3000, RRID:AB_2247211, Cell Signaling Technology, USA), mouse anti-NeuN (1:800, RRID:AB_2313673, Millipore, USA), mouse anti-CaMKII α (1:100, RRID:AB_626789, Santa Cruz Biotechnology, USA), mouse anti-CaMKII α (1:50, RRID:AB_2721906,Cell Signaling Technology, USA), Rabbit anti-ADRB1 (1:100, RRID:AB_10885544, Bioss, China), followed by the corresponding fluorophore-conjugated secondary antibodies for 1.5 h at room temperature. The following secondary antibodies were used here, including Alexa Fluor 555-labeled donkey anti-rabbit secondary antibody (1:500, RRID: AB_2762834, Thermo Fisher Scientific, USA), Alexa Fluor 488-labeled donkey anti-mouse secondary antibody (1:500, RRID: AB_141607, Thermo Fisher Scientific, USA). In detail of Figure 1k-l, brain slices were attached to slides and placed in an oven at 60°C for 40 min, and then immersed in 4% PFA solution for 15 min in a refrigerator at 4°C for fixation. Then graded concentration alcohol dehydration (50%-70%-100%-100%) was performed sequentially for 5 min each time. Next, antigen repair was performed by immersing the slides with brain slices attached in 10 mM PH=6.0 trisodium citrate solution and heating in a water bath at 82°C for 30 min. Normal staining steps were then followed. Fluorescence signals were visualized using a Leica DM6B upright digital research microscope (Leica, Germany) and Leica TCS SP8 confocal microscope (Leica, Germany).

**Figure 1.**
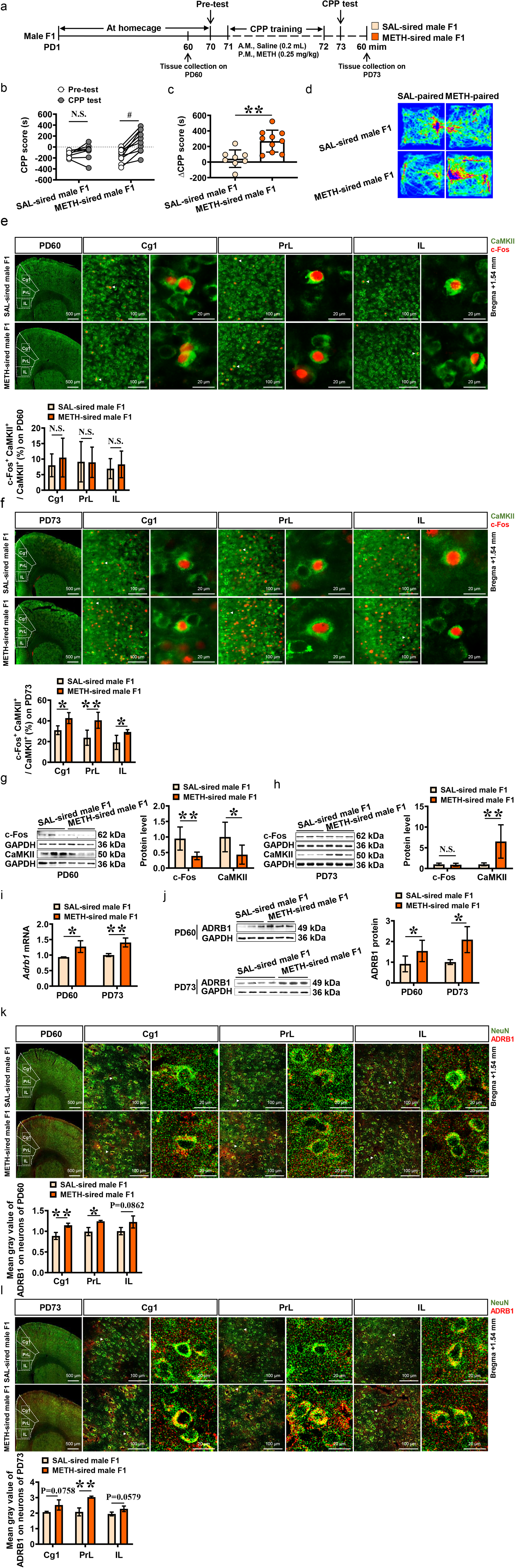
The CPP, mPFC activity and ADRB1 level in male F1. **a**, Experimental design and timeline. **b**, CPP score by male F1. **c**, ΔCPP score by male F1. **d**, Traveled traces in CPP apparatus by male F1. **e-f**, The percentage of c-Fos-positive cells in Cg1, PrL and IL on PD60 and on PD73. CaMKII and c-Fos were used as markers for the pyramidal neurons and activated neurons, respectively. Scale bar, 500 μm /100 μm /20 μm. **g-h**, Levels of c-Fos and CaMKII protein on PD60 and PD73. **i**, Levels of *Adrb1* mRNA on PD60 and PD73. **j**, Levels of ADRB1 protein on PD60 and PD73. **k-l**, Mean gray value of ADRB1 on mPFC neurons on PD60 and PD73. Scale bar, 500 μm /100 μm /20 μm. Cg1, cingulate cortex; PrL, prelimbic cortex; IL, infralimbic cortex. SAL-sired male F1, saline-sired male F1 mice; METH-sired male F1, methamphetamine-sired male F1 mice. N.S., *P* > 0.05. #, *P* < 0.05 vs baseline (Pre-test CPP score). *, *P* < 0.05, **, *P* < 0.01 vs SAL-sired male F1.

### Western blot

Total protein was extracted from mPFC of individual mouse using RIPA lysis buffer (Beijing ComWin Biotech Co., China), Phosphatase Inhibitor (Beijing ComWin Biotech Co., China), and Protease Inhibitor (Beijing ComWin Biotech Co., China) according to the manufacturer’s instructions. Protein samples (15 or 20 μg) was separated by 10% SDS–PAGE, 12% SDS–PAGE and electrophoretically transferred onto PVDF membranes. The transferred membranes were blocked with 5% non-fat dry milk and 0.1% Tween 20 in TBST buffer for 2 h at room temperature, then subsequently incubated with the following primary antibodies: Rabbit anti-c-Fos (1:1000, RRID:AB_224721, Cell Signaling Technology, USA), Rabbit anti-GAPDH (1:10000, RRID:AB_2651132, Bioworld Technology, USA), Rabbit anti-CaMKII α (1:1000, RRID:AB_305050, Abcam, USA), Rabbit anti-ADRB1 (1:1000, RRID:AB_10885544, Bioss, China), Rabbit anti-ARRB2 (1:1000, BA3767, Boster, China), Rabbit anti-ERK1/2 (1:1000, BM4326, Boster, China), Rabbit anti-phospho-ERK1/2 (1:1000, BM4156, Boster, China), Rabbit anti-ΔFosB (1:1000, bsm-52071R, Bioss, China), Rabbit anti-PKA (1:1000, bs-0520R, Bioss, China), Rabbit anti-CREB (1:1000, RRID:AB_2800317, Cell Signaling Technology, USA), Rabbit anti-phospho-CREB (1:1000, RRID:AB_2561044, Cell Signaling Technology, USA). The next day, the membranes were washed in Tris-buffered saline with Tween 20 (TBST) and incubated with horseradish peroxidase (HRP)-conjugated secondary antibody goat anti-rabbit or goat anti mouse (1:5000, Beijing ComWin Biotech Co., China) at room temperature for 1 h. The blots were visualized by the eECL Western Blot kit (Beijing ComWin Biotech Co., China, CW0049S) or ELC Kit (Vazyme^™^) and the signal was visualized by imaging system (Tanon-5200, Shanghai, China). The blots were washed with stripping buffer (Beyotime Institute of Biotechnology, China) to be reprobed with other antibodies. In this study, GAPDH was used as the loading control. The relative level of each protein expression was normalized to GAPDH. Values for target protein levels were calculated using Image J software (NIH, USA).

### RNA extraction and quantitative real-time PCR

The mPFC were dissected immediately on ice. Total RNA was extracted from the mPFC using FastPure Cell/Tissue Total RNA Isolation Kit (Vazyme, China), according to the manufacturer’s instructions. All samples were tested for concentration and purified using a 4200 TapeStation (Agilent Technologies, Santa Clara, USA). After RNA extraction, the amount of RNA was normalized across samples, and cDNA was created using HiScript II Q RT SuperMix for qPCR (+gDNA wiper) (Vazyme, China). *Gapdh* was used as the internal control. The primers used in the study were as follows: *Adrb1* (forward, CTCATCGTGGTGGGTAACGTG; reverse, ACACACAGCACATCTACCGAA). *c-Fos* (forward, CGGGTTTCAACGCCGACTA; reverse, TTGGCACTAGAGACGGACAGA). *Gapdh* (forward, AGGTCGGTGTGAACGGATTTG; reverse, TGTAGACCATGTAGTTGAGGTCA). The *Adrb1* and *c-Fos* mRNA levels were normalized to *Gapdh* mRNA levels. The relative mRNA level was calculated by the comparative CT method (2^-ΔΔCt^).

### Local administration of betaxolol in mPFC

The cannula-double (O.D. 0.41 mm - 27G, C.C 0.8 mm, Mates with M 3.5, Cat 62063, RWD, China) were planted into mPFC (AP +2.7 mm, ML ± 0.4 mm, DV – 1.5 mm) of male F1 mice on PD63 before CPP training. After surgery, mice were maintained at home cage about 1 week. From PD70, male F1 mice were subjected to METH-induced CPP training. In Experiment 1 and S1, a volume of 500 nL of betaxolol (specific antagonist for ADRB1) were bilaterally injected into mPFC at a rate of 100 nL/min at 15 min prior to CPP test on PD73. In Experiment 2 and S2, a volume of 500 nL of betaxolol bilaterally into the mPFC at a rate of 100 nL/min at 15 min prior to CPP training on PD71 and PD72.

### Specific knockdown of ADRB1 on mPFC CaMKII-positive neurons

On PD55, mice were anesthetized with 2% isoflurane in oxygen, and were fixed in a stereotactic frame (RWD, Shenzhen, China). A heating pad was used to maintain the core body temperature of the mice at 36°C. The coordinates of mPFC were defined as AP + 2.05 mm, ML ± 0.3 mm, DV – 1.9 mm. A volume of 400 nL virus of *CaMKIIap-EGFP-MIR155 (MCS)-SV40PolyA* (KD group) (Titer, 2.41F + 13 v.g/mL; Cat, GIDV0286541, GeneChem, Shanghai, China) or *CaMKIIap-EGFP-SV40PolyA* (Ctrl group) (Titer, 4.01E + 12 v.g/mL; Cat, AAV9/CON540, GeneChem, Shanghai, China) were injected bilaterally into mPFC at a rate of 80 nL/min. After surgery, mice were kept at homecage about 3 weeks.

### Dendritic spine analysis

The dendritic spines of mPFC were photographed using the z-stack scanning function of Leica TCS SP8 confocal microscope (Leica, Germany). The Spine quantification were performed under LAS X software (Leica, Germany). The number of spines is counting only in the second or the third branches of dendrites from the body of CaMKII-positive neurons. Density of spines were scored in about 50 μm dendritic segments in length.

### Statistical analysis

Statistical analysis was carried out using GraphPad Prism 8.0.2 software. All data are presented as the Mean ± SD. CPP score are analyzed by two-way analysis of variance (ANOVA) with Sidak’s multiple comparisons test, and other data are analyzed by unpaired *t*-tests. Statistical significance was set as *P* < 0.05.

## RESULTS

### METH-sired male F1 mice acquires METH^F1^-induced CPP in adulthood

In the present study, the 0.25 mg / kg dose of METH administration (METH^F1^), being insufficient to produce CPP in naïve mice (subthreshold dose), was used to induce CPP in male F1 mice (Figure 1a) and female F1 mice (Figure S1a).

In male F1 mice (Figure 1b-d), METH^F1^ failed to induce CPP in SAL-sired male F1 (n = 8 mice, *t* = 0.9474, *P* = 0.5872, Figure 1b), but increased CPP score of METH-sired male F1 (n = 10 mice, *t* = 6.587, *P* < 0.0001, Figure 1b) when compared with the corresponding Pre-test data. The ΔCPP score (*t* = 3.685, *P* = 0.002, Figure 1c) in METH-sired male F1 was higher than that in SAL-sired male F1.

In female F1 mice (Figure S1b-d), METH^F1^ failed to induce CPP in both SAL-sired female F1 (n = 8 mice, *t* = 1.398, *P* = 0.3295, Figure S1b) and METH-sired female F1 (n = 10 mice, *t* = 1.979, *P* = 0.1264, Figure S1b) when compared with the corresponding Pre-test data. There was no significance on ΔCPP score (*t* = 0.277, *P* = 0.7854, Figure S1c) between METH-sired and SAL-sired female F1.

These results indicate that paternal METH exposure induce higher sensitivity to METH in male, but not female offspring. As such, only male F1 mice were used to the subsequent experiments.

### METH^F1^ activates more CaMKII-positive neurons and induces higher ADRB1 levels in mPFC of METH-sired male F1 mice

The mPFC tissue were collected from male F1 mice on PD60 and PD73, which represent the baseline condition and responsive condition to METH^F1^, respectively. The c-Fos, CaMKII and NeuN are used as markers to label neuronal activation, glutamatergic neuron and total neuron in mPFC, respectively.

On PD60, there was no significance in the percentage of c-Fos-positive & CaMKII-positive neurons in subregions of mPFC including cingulate cortex (Cg1, *t* = 0.595, *P* = 0.5839), prelimbic cortex (PrL, *t* = 0.0371, *P* = 0.9722) and infralimbic cortex (IL, *t* = 0.4451, *P* = 0.6793) between METH-sired male F1 (n = 3 mice) and SAL-sired male F1 (n = 3 mice, Figure 1e). While on PD73 following METH^F1^, the percentage of c-Fos-positive & CaMKII-positive neurons were at higher levels in Cg1 (*t* = 3.447, *P* = 0.0137), PrL (*t* = 3.539, *P* = 0.0076) and IL (*t* = 3.160, *P* = 0.0134) of METH-sired male F1 (n = 4-5 mice) than that of SAL-sired male F1 (n = 4-5 mice, Figure 1f). On PD73, the neurons (labeled as NeuN-positive neurons) were more activated in Cg1 (*t* = 3.031, *P* = 0.0163), PrL (*t* = 4.227, *P* = 0.0022) and IL (*t* = 4.098, *P* = 0.0027, Figure S2a), and the dendritic spine density of CaMKII-positive neurons was increased (*t* = 4.960, *P* = 0.0026, Figure S2b) in mPFC of METH-sired male F1 (n = 4-6 mice) than of SAL-sired male F1 (n = 4-6 mice). As one of important kinase, CaMKII can also be used to reflect the activities of neurons. On PD60, the baseline levels of mPFC c-Fos (*t* = 3.496, *P* = 0.0058) and CaMKII protein (*t* = 2.457, *P* = 0.0338) were obviously lower in METH-sired male F1 (n = 6 mice) than that in SAL-sired male F1 (n = 6 mice, Figure 1g). On PD73 following METH^F1^, there was similar levels of c-Fos protein (*t* = 0.5061, *P* = 0.6238) but a higher levels of CaMKII protein (*t* = 3.350, *P* = 0.0074) in METH-sired male F1 (n = 6 mice) than in SAL-sired male F1 (n = 6 mice, Figure 1h). These results indicate that paternal METH exposure did not change or even reduce the mPFC activity, but obviously increased mPFC response to METH^F1^, especially the CaMKII-positive neurons in it.

On PD60, the baseline levels of mPFC *Adrb1* gene (*t* = 3.084, *P* = 0.0368, Figure 1i) and ADRB1 protein (*t* = 2.400, *P* = 0.0373, Figure 1j) were higher in METH-sired male F1 (qPCR: n = 3 mice; protein: n = 6 mice) than that in SAL-sired male F1 (qPCR: n = 3 mice; protein: n = 6 mice). On PD73 following METH^F1^, mPFC *Adrb1* gene (*t* = 4.621, *P* = 0.0099, Figure 1i) and ADRB1 protein (*t* = 3.489, *P* = 0.0175, Figure 1j) were persistently at higher levels in METH-sired male mice (qPCR: n = 3 mice; protein: n = 3 mice) than that in SAL-sired male F1 (qPCR: n = 3 mice; protein: n = 4 mice).

Further, ADRB1 levels on neurons of mPFC and its subregions (Figure 1k-l) were examined by immunostaining. On PD60, the mean gray value of ADRB1 on neurons was much higher in the Cg1 (*t* = 4.649, *P* = 0.0097) and PrL (*t* = 4.311, *P* = 0.0125), and a high trend in IL (*t* = 2.265, *P* = 0.0862) of METH-sired male F1 (n = 3 mice) than that in SAL-sired male F1 (n = 3 mice). On PD73 following METH^F1^, the mean gray value of ADRB1 on neurons was continuously higher in PrL (*t* = 6.304, *P* = 0.0032), and a higher trend in Cg1 (*t* = 2.382, *P* = 0.0758), IL (*t* = 2.635, *P* = 0.0579) of METH-sired male F1 (n = 3 mice), when compared to that in SAL-sired male F1 (n = 3 mice).

To observe the effects of METH^F1^ treatment on levels of c-Fos and ADRB1 on the mPFC under normal conditions, male naïve mice were subjected to subthreshold dose of METH treatment (Figure S3a). Neither *c-Fos* gene (*t* = 0.4522, *P* = 0.6745, Figure S3b) and *Adrb1 (t* = 0.9778, *P* = 0.3835, Figure S3b), nor c-Fos protein (*t* = 1.115, *P* = 0.3274, Figure S3c) and ADRB1 protein (*t* = 1.723, *P* = 0.1600, Figure S3c) in mPFC were changed by subthreshold dose of METH administration in naïve mice (n = 3 mice per group).

With these results, we infer that paternal METH exposure-increased ADRB1 on mPFC neurons might contribute to the higher sensitivity to METH in male F1 mice.

### Blocking ADRB1 suppresses METH^F1^-induced CPP in METH-sired male F1 mice

Here, betaxolol was bilaterally injected into mPFC (Figure S4a) to block the activation of ADRB1. In Experiment 1 (Figure 2a), a single dose of betaxolol were injected into mPFC at 15 min prior to METH^F1^-induced CPP test. METH-sired male F1 did not exhibit METH-preferred behavior any more (n = 8 mice, *t* = 1.666, *P* = 0.2219), and METH-sired male F1 with vehicle treatment exhibited METH^F1^-induced CPP continuously (n = 8 mice, *t* = 3.518, *P* = 0.0068). The ΔCPP score (*t* = 3.666, *P* = 0.0025) in vehicle-injected METH-sired male F1 was much higher than that in betaxolol-injected METH-sired male F1 (Figure 2b-d).

**Figure 2.**
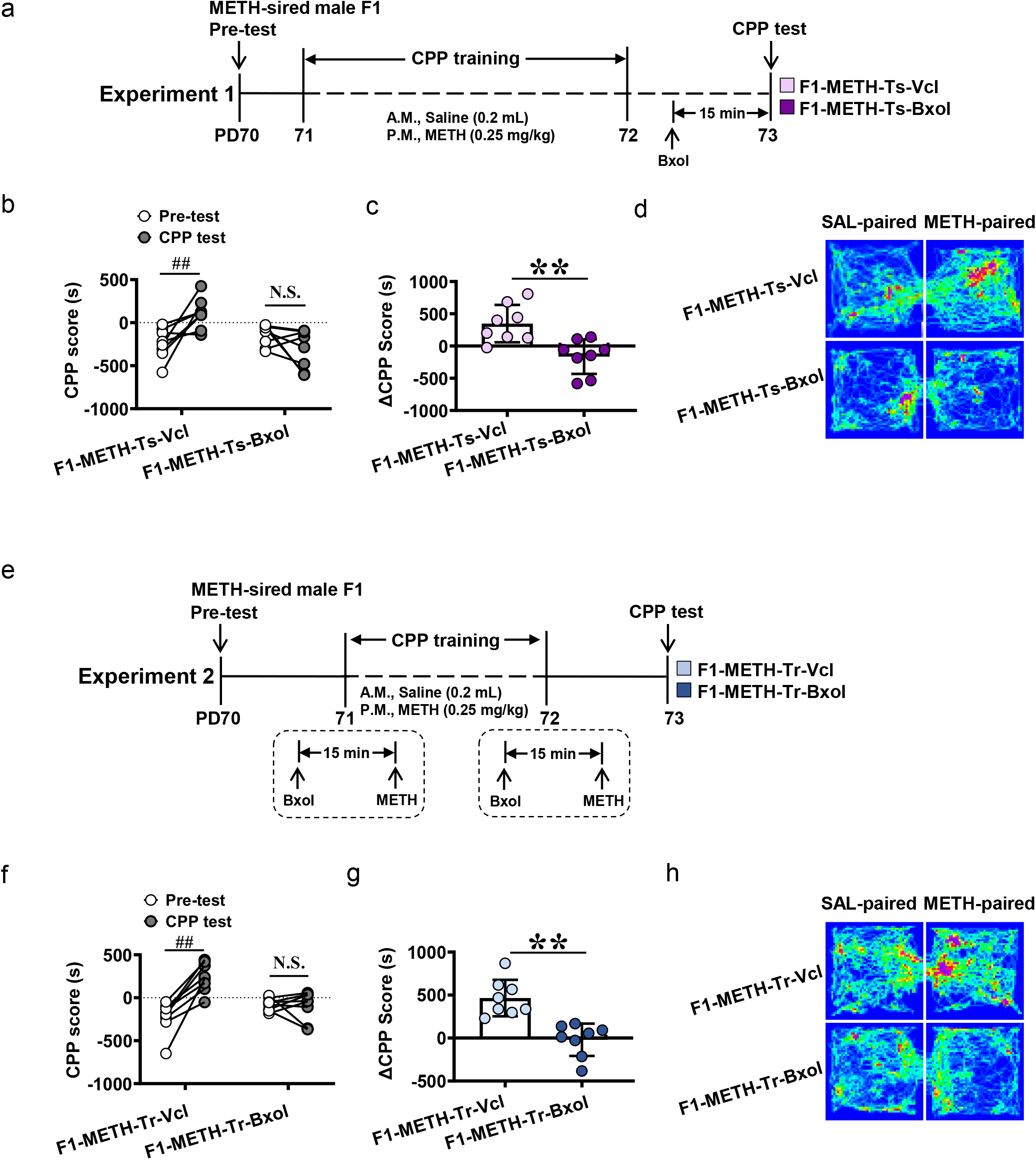
Effects of betaxolol on METH^F1^-induced CPP in METH-sired male F1. **a**, Timeline of Experiment 1. **b**, CPP score. **c**, ΔCPP score. **d**,Traveled traces in the CPP apparatus by METH-sired male F1. **e**, Timeline of Experiment 2. **f**, CPP score. **g**, ΔCPP score. **h**, Traveled traces in the CPP apparatus by METH-sired male F1. F1-METH-Ts-Vcl, vehicle bilaterally into the mPFC of METH-sired male F1 at 15 min prior to CPP test; F1-METH-Ts-Bxol, betaxolol bilaterally into the mPFC of METH-sired male F1 at 15 min prior to CPP test; F1-METH-Tr-Vcl, vehicle bilaterally into the mPFC of METH-sired male F1 during CPP training; F1-METH-Tr-Bxol, betaxolol bilaterally into the mPFC of METH-sired male F1 during CPP training. N.S., *P* > 0.05. ##, *P* < 0.01 vs Pre-test CPP score. **, *P* < 0.01 vs F1-METH-Ts-Vcl or F1-METH-Tr-Vcl.

In Experiment 2 (Figure 2e), betaxolol were injected into mPFC during the whole period of CPP training. Betaxolol-treated METH-sired male F1 mice did not acquire METH^F1^-induced CPP (n = 8 mice, *t* = 0.2852, *P* = 0.9514), but vehicle-treated METH-sired male F1 mice still exhibit METH^F1^-induced CPP (n = 8 mice, *t* = 6.570, *P* < 0.0001). The ΔCPP score (*t* = 4.847, *P* = 0.0003) in vehicle-injected METH-sired male F1 was much higher than that in betaxolol-injected METH-sired male F1 (Figure 2f-h).

Here, we also examine the effects of betaxolol on METH^F1^-induced CPP in SAL-sired male F1 mice. In Experiment S1 (Figure S4b-e), similar to the above results, METH^F1^ could not induce CPP in SAL-sired male F1 (n = 5 mice, *t* = 0.1464, *P* = 0.9872). Further, a single dose of betaxolol administration at 15 min prior to CPP did not affect their CPP score as well (n = 6 mice, *t* = 0.1320, *P* = 0.9896). There was no difference in the ΔCPP score (*t* = 0.01917, *P* = 0.9851) between vehicle-treated and betaxolol-treated SAL-sired male F1. In Experiment S2 (Figure S4f-i), similarly, METH^F1^ treatment could not induce CPP in SAL-sired male F1 (n = 6 mice, *t* = 0.5901, *P* = 0.8136). The betaxolol injection during CPP training did not significantly affect their CPP scores as well (n = 6 mice, *t* = 0.4447, *P* = 0.8884). There was no difference in the ΔCPP score (*t* = 0.4406, *P* = 0.6689) between vehicle-treated and betaxolol-treated SAL-sired male F1.

These results indicate that mPFC ADRB1 mediate both formation (CPP training) and expression (CPP test) of METH^F1^-induced CPP in METH-sired F1 mice, and contribute to paternal METH exposure-increased sensitivity to METH in male F1 mice.

### Knocking-down ADRB1^CaMKII^ blocks METH-induced CPP in METH-sired male F1 mice

Here, adeno-associated virus (AAV) were used to knocking-down ADRB1 on CaMKII-positive neurons (ADRB1^CaMKII^) within mPFC (Figure 3a). As shown in Figure 3b, AAV of *CaMKIIap-EGFP-MIR155(MCS)-SV40PolyA* (KD group) or AAV of *CaMKIIap-EGFP-SV40PolyA* (Ctrl group) were bilaterally infused into mPFC on P55, which could specifically transfect CaMKII-positive neurons. On P70, about 60%~80% CaMKII-positive neurons were transfected with virus (Figure 3c). The levels of *Adrb1* gene (*t* = 7.361, *P* = 0.0018, Figure 3d), ADRB1 protein (*t* = 4.286, *P* = 0.0128, Figure 3e), and ADRB1^CaMKII^ expression (*t* = 4.237, *P* = 0.0133, Figure 3f) were all reduced in mPFC of male F1 by KD virus (n = 3 mice), when compared with Ctrl virus (n = 3 mice), indicating good efficiency of KD virus.

**Figure 3.**
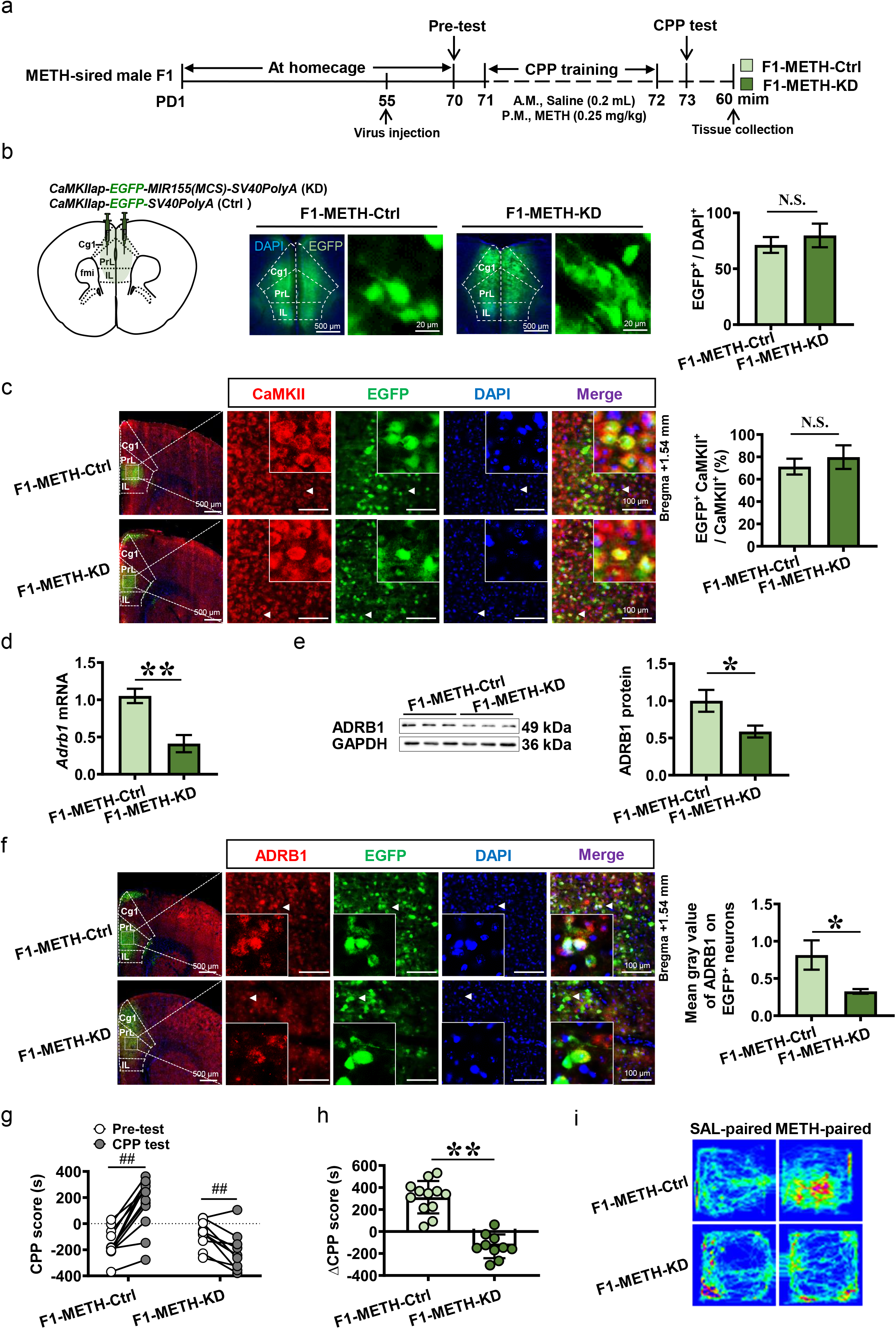
Effects of knocking-down ADRB1^CaMKII^ on METH^F1^-induced CPP in METH-sired male F1. **a**, Experimental design and timeline. **b**, Schematic diagram of viral injection and representative images of viral transfection in mPFC (middle pannel). Scale bar, 500 μm /20 μm. **c**, Percentage of viral transfected neurons in the total pyramidal neurons of mPFC. Scale bar, 500 μm /100 μm. **d**, Levels of *Adrb1* mRNA in mPFC. **e**, Levels of ADRB1 protein in mPFC. **f**, Mean gray value of ADRB1 on virus transfected neurons of mPFC. Scale bar, 500 μm /100 μm. **g**, CPP score. **h**, .ΔCPP score. **i**, Traveled traces in the CPP apparatus by METH-sired male mice. F1-METH-Ctrl, METH-sired male F1 mice injected with Ctrl virus. F1-METH-KD, METH-sired male F1 mice injected with KD virus. N.S., *P* > 0.05. ##, *P* < 0.01 vs pre-test CPP score. *, *P* < 0.05, **, *P* < 0.01 vs F1-METH-Ctrl.

As shown Figure 3g-i, Ctrl virus-treated METH-sired male F1 mice still acquired METH^F1^-induced CPP in (n = 12 mice, *t* = 8.228, *P* < 0.0001), but KD virus-treated METH-sired male F1 mice did not exhibit METH-preferred behaviors and even produced METH-aversion behaviors (n = 10 mice, *t* = 3.253, *P* = 0.008), when compared with their corresponding Pre-test score. The ΔCPP score of KD group was much lower than that in Ctrl group (*t* = 7.95, *P* < 0.0001).

These results confirmed that mPFC ADRB1^CaMKII^ mediate paternal METH exposure-induced higher sensitivity to METH in male F1 mice.

### Knocking-down ADRB1^CaMKII^ reduces mPFC activation in METH-sired male F1 mice

Compared to Ctrl group (protein: n = 3 mice; immunostaining: n = 6 mice; dendritic spine density: n = 6 mice), knocking-down mPFC ADRB1^CaMKII^ obviously decreased levels of c-Fos protein (*t* = 5.372, *P* = 0.0058, Figure 4a) and immunostaining (*t* = 3.136, *P* = 0.0106, Figure 4b), and dendritic spine density of CaMKII-positive neurons (*t* = 3.891, *P* = 0.003, Figure 4c) in mPFC of METH-sired male F1 (protein: n = 3 mice; immunostaining: n = 6 mice; dendritic spine density: n = 6 mice).

**Figure 4.**
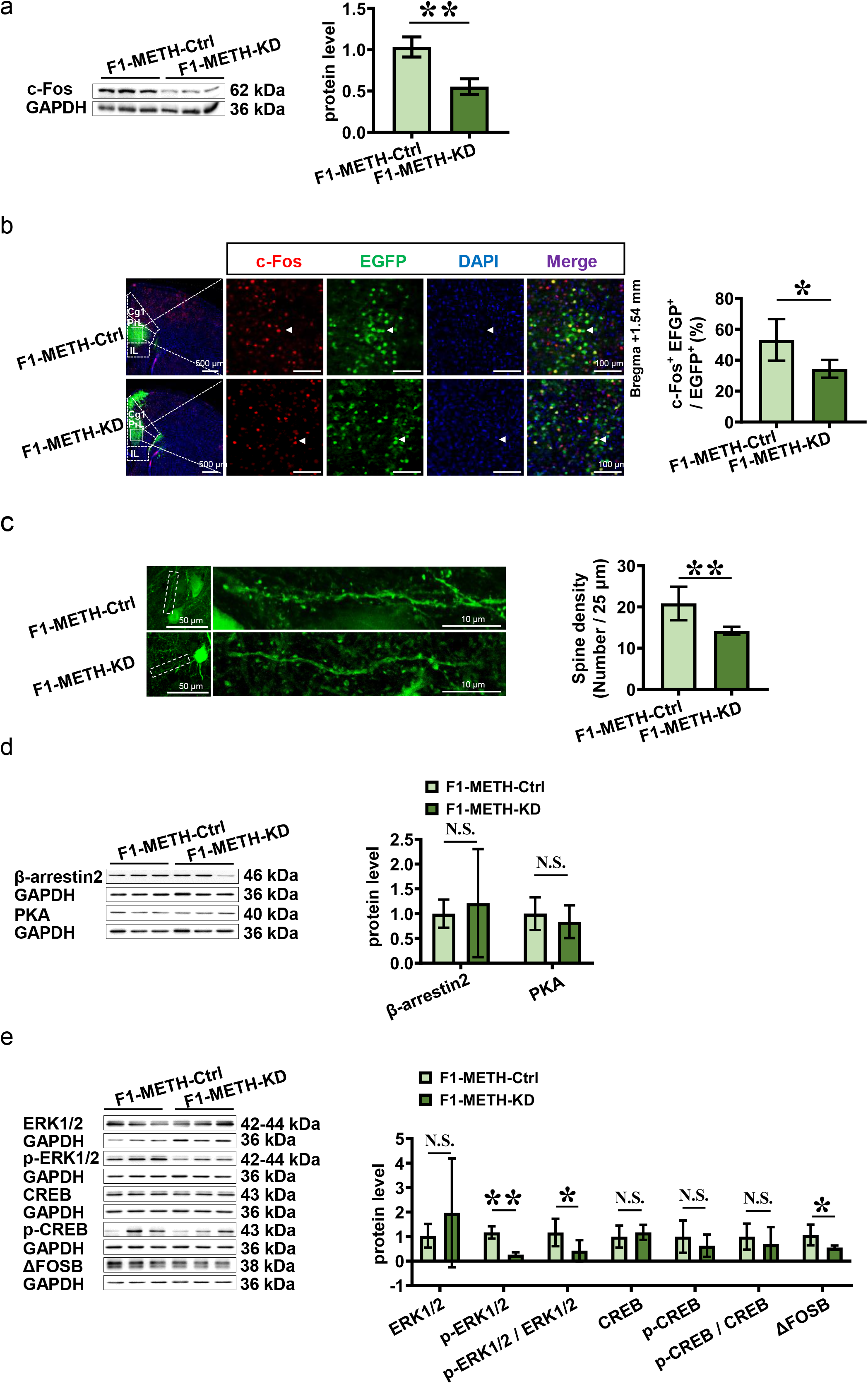
Effects of knocking-down ADRB1^CaMKII^ on mPFC activity and subsequent signals in METH-sired male F1. **a**, Levels of c-Fos protein. **b**, The c-Fos immunostaining. Scale bar, 500 μm /100 μm. **c**, Density of dendritic spine. Scale bar, 50 μm /10 μm. **d**, Levels of β-arestin2 and PKA protein. **e**, Levels of ERK1/2, p-ERK1/2, ERK1/2/p-ERK1/2, CREB, p-CREB, p-CREB/ CREB and ΔFosB protein. F1-METH-Ctrl, METH-sired male F1 mice injected with Ctrl virus. F1-METH-KD, METH-sired male F1 mice injected with KD virus. N.S., *P* > 0.05. *, *P* < 0.05, **, *P* < 0.01, vs F1-METH-Ctrl.

The potential subsequent molecules of β-adrenergic signaling pathways especially β-arrestin2 and PKA, were examined in mPFC. Compared to Ctrl virus group (n = 3-6 mice), protein levels of β-arrestin2 (*t* = 0.4597, *P* = 0.6555) and PKA (*t* = 0.6055, *P* = 0.5775, Figure 4d) were not changed, the p-ERK1/2 (*t* = 8.202, *P* < 0.0001), ratio of p-ERK1/2 to ERK1/2 (*t* = 2.486, *P* = 0.0347) and ΔFosB (*t* = 2.944, *P* = 0.0147) were decreased, and ERK1/2 (*t* = 0.9153, *P* = 0.3839), CREB (*t* = 0.5454, *P* = 0.6145), p-CREB (*t* = 0.8085, *P* = 0.4641) and ratio of p-CREB to CREB (*t* = 0.6042, *P* = 0.5783) were not changed in mPFC by knocking-down ADRB1 (n = 3-6 mice, Figure 4e).

These results indicate that ADRB1^CaMKII^ positively activate mPFC CaMKII-positive neurons, p-ERK1/2 and ΔFosB are the subsequent signals for ADRB1 in the process.

## DISCUSSION

The current study found that paternal METH exposure induces higher ADRB1 levels on mPFC neurons. The subthreshold dose of METH (METH^F1^), which is insufficient to produce CPP normally, efficiently induced METH-related CPP in METH-sired male F1, accompanied with more activated CaMKII-positive neurons and continuous higher ADRB1 in their mPFC. Either inhibiting ADRB1 function or knocking-down ADRB1 expression on CaMKII-positive neurons (ADRB1^CaMKII^) efficiently reduced METH^F1^-evoked mPFC activation and blocked METH^F1^-induced CPP in METH-sired male F1. These results indicate that paternal METH exposure induce a higher sensitivity to METH in male offspring potentially through driving mPFC ADRB1^CaMKII^ (Figure 5).

**Figure 5.**
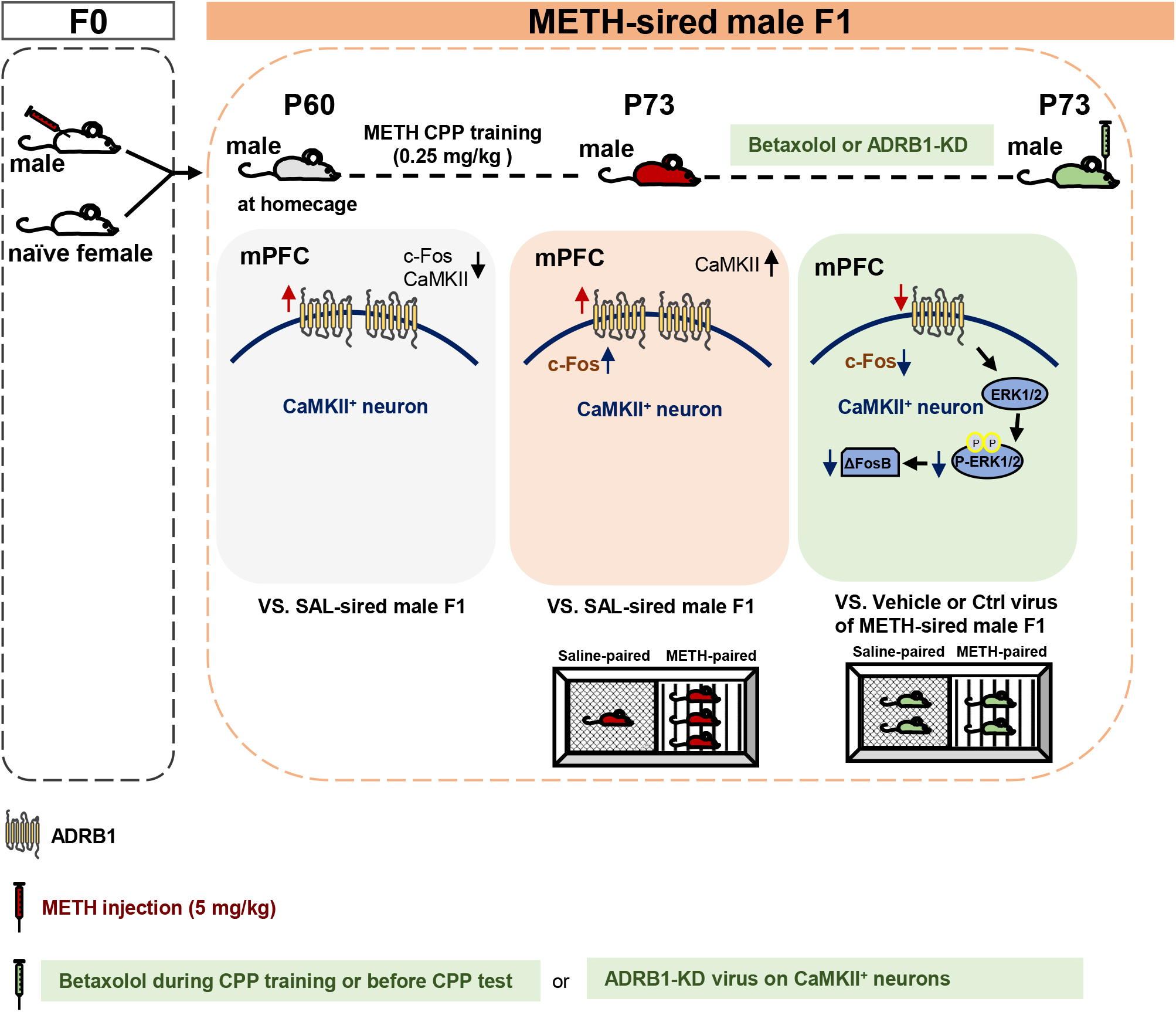
Schematic summary of this study. Paternal methamphetamine (METH) exposure induces higher ADRB1 levels on mPFC neurons in male F1 mice during adulthood. Subthreshold dose of METH treatment in F1 (METH^F1^), which is not sufficient to induce CPP, efficiently produces METH-related CPP in METH-sired male F1 mice, accompanied with more activated mPFC CaMKII-positive neurons than that in saline-sired male F1 mice. Locally inhibiting ADRB1 function or specifically knocking-down ADRB1 on CaMKII-positive neurons (ADRB1^CaMKII^) efficiently reduces METH^F1^-evoked mPFC activation, and ultimately blocks METH^F1^-induced CPP in METH-sired male F1 mice. The p-EKR1/2 and ΔFosB might be potential subsequent signals of ADRB1 in the process.

METH is commonly abused psychostimulant among men of reproductive age. Consistent with previous studies, we found that METH^F1^, which is insufficient to induce CPP in naïve mice, effectively induced METH-preferred behaviors in METH-sired male F1 and not in SAL-sired male F1, indicating an enhanced sensitivities to drugs in F1 of sires exposed to METH. The mPFC has been implicated in the vulnerability to drug abuse [6–8]. Currently, little literature uncover the association of paternal METH intake with offspring mPFC alteration. *In utero* exposure to cocaine had been reported to increase the induction of long-term potentiation (LTP) and neuronal excitability at pyramidal neurons in mPFC of offspring [29], an increase in dendritic spines of II/III layer of pyramidal neurons in mPFC of offspring [30], and a reduction of the sensitivity of locomotor activity to cocaine in offspring rats [29, 31]. By contrast, Yan et al. [32] reported prenatal cocaine exposure attenuated baseline levels of mature BDNF protein in offspring brain, and Cantacorps et al. [33] found prenatal alcohol exposure increased vulnerability to cocaine addiction in adult mice via a greater glutamate receptor in mPFC. Similarly, human imaging studies demonstrated that prenatal cocaine exposed-adolescents exhibited less capability of increasing their PFC activation in response to memory load [34], and infants with prenatal cocaine use had a lesser gray matter in prefrontal brain regions [35, 36]. In addition, adolescents with prenatal cocaine exposure had reduced activity in PFC when exposing to favorite-food cues [37]. Although there are conflicting results among these studies, they suggest that plasticity of frontal brain play essential roles in prenatal drugs intaking-elicited heritable susceptibility to addiction in offspring. Here, we found that paternal METH exposure seemed to reduce mPFC activity in male F1, as indicated by lower levels of c-Fos and CaMKII in mPFC. However, METH^F1^ treatment, which could not activate mPFC in naïve mice, reversely triggered mPFC activation in METH-sired male F1, implying potential intrinsic molecules in mPFC that may contribute to paternal METH exposure-increased sensitivity to METH in male F1 mice.

ADRB1 is highly expressed on glutamatergic neurons of mPFC. In general, endogenous activation of ADRB1 excites neurons via eliciting the depolarization on the membrane potential in mPFC [38]. An early study found that prenatal cocaine exposure increased cortical ADRB1 about 68% in pup rats at 30 days of age, a time when these rats showed hyperactivity [39]. Unfortunately, few studies further explored the role of this increase of cortical ADRB1 in addiction or related psychiatric disorders of offspring. Administration of betaxolol, a selective ADRB1 antagonist, blocked reinstatement of cocaine CPP [19]. Further, blocking ADRB1 impaired reconsolidation and reinstatement of cocaine self-administration [40]. Here, we found that paternal METH exposure increased ADRB1 on mPFC neurons. We believe that this higher ADRB1 is kind of compensatory effect for brain injury in offspring of sires exposed to drugs, which could strengthen the sensitivity of neurons to internal or external stimuli. Following METH^F1^ treatment, we observed a continuous higher ADRB1 level in METH-sired male F1. Blocking ADRB1 or knocking-down ADRB1 on CaMKII-positive neurons (ADRB1^CaMKII^) efficiently reduced METH^F1^-evoked mPFC activation, and ultimately blocked METH^F1^-induced CPP in METH-sired male F1. There results indicate that ADRB1^CaMKII^ mediate mPFC activity, which is one key intrinsic molecule of mPFC for the paternal METH exposure-induced higher sensitivity in male offspring.

ADRB1 activates neurons by eliciting its downstream protein kinases and signal pathways (e.g. PKA, β-arrestin2), or mediating the function of nearby channels (e.g. TREK-2-like channels) or receptors (e.g. GluR1) on the membrane of cells. The critical role of ADRB1/cyclic-AMP dependent PKA in synaptic plasticity of the frontal cortex are well-studied [41]. Recently, Huang et al. found that ADRB1/β-arrestin/ERK signaling in entorhinal cortex, acting as a non-classical G protein-coupled pathway, mediated memory reconsolidation [28]. Their lab later reported that overexpression of β-arrestin2 in PFC promoted extinction learning of cocaine-induced CPP, which was blocked by propranolol, a nonbiased β-adrenergic receptor antagonist [42]. Thus, we detected the molecules related to ADRB1, especially those of β-arrestin2-dependent and PKA-dependent signal proteins. Unexpected, the levels of β-arrestin2 and PKA failed to change in mPFC after knocking-down ADRB1. While, we found that knocking-down ADRB1 effectively reduce the levels of p-ERK1/2 and ΔFosB in mPFC. Son et al. [43] has been reported that acupuncture exert its therapeutic effects on METH-induced addiction via suppression of METH-increased p-ERK1/2 and ΔFosB in PFC. The p-ERK1/2 signaling cascades in PFC has been thought of important contributor for the development of METH addiction [44]. ΔFosB, a transcription factor, is increased in PFC by drugs including METH [43] and Δ9-tetrahydrocannabinol (THC) [45], which leads to a long-lasting enhanced behavioral response.

Further, ΔFosB is preferentially induced by cocaine in adolescent animals, which engaged in the higher vulnerability to addiction [46]. Unlike c-Fos, Arc and ERK, ΔFosB maintains adaptive changes in brain for a long time following drugs of abuse [47], which is involved in the long-term pathological plasticity of brain and persistent addictive behavior by drugs [48]. Together with our present study, these findings revealed that p-ERK1/2 and ΔFosB may be subsequent signals for ADRB1 mediating heritable susceptibility to addiction in offspring.

Collectively, our findings reveal a novel molecular mechanism underlying paternal METH exposure-induced higher sensitivity to addiction in male offspring, and mPFC ADRB1^CaMKII^ is a promising pharmacological target for predicting or treating transgenerational susceptibility to addiction, especially caused by paternal drug exposure.

## ACKNOWLEDGMENTS

This work is supported by National Natural Science Foundation of China (82271531 and 82071495), Natural Science Foundation of Jiangsu Province, China (BK20201398) and Natural Science Foundation of the Higher Education Institutions of Jiangsu Province, China (21KJB360007).

## AUTHOR CONTRIBUTIONS

Zheng Y and Liu D have equal contribution to the manuscript. Zheng Y, Liu D and Li Z performed behavioral tests, morphological tests. Guo H, Chen W and Liu Z performed molecular experiments. Hu T, Zhang Y, Li X, Fan Y carried out the virus-related experiment. Fan Y and Liu D performed data analysis. Zhao Z, Cai Q and Ge F assist the data analysis. Guan X and Fan Y wrote the manuscript. Liu D help to support the proposal. Guan X developed the overall concept.

## DATA AVAILABILITY STATEMENT

The data that support the findings of this study are available from the corresponding author upon reasonable request.

