## Supplementary material for "Paternal methamphetamine exposure induces higher sensitivity to methamphetamine in male offspring through driving ADRB1 on CaMKII-positive neurons in mPFC": Suppl legends and figures

**Supplementary Data**

**Supplementary Figure Legends**

**Figure S1. METH-induced CPP in female F1. a**, Experimental design and timeline. **b**, CPP score by female F1. **c**, ΔCPP score by female F1. **d,** Traveled traces in CPP apparatus by female F1. SAL-sired female F1, saline-sired female F1 mice; METH-sired female F1, methamphetamine-sired female F1 mice. N.S., $P > 0.05$ vs Pre-test CPP score or SAL-sired female F1.

**Figure S2. The mPFC activity in male F1. a,** The number of c-Fos-positive neurons. Scale bar: 500 µm /100 µm. **b,** Density of dendritic spine. Scale bar: 50 µm /10 µm. Cg1, cingulate cortex; PrL, prelimbic cortex; IL, infralimbic cortex. SAL-sired male F1, saline-sired male F1 mice; METH-sired male F1, methamphetamine-sired male F1 mice. *, $P < 0.05$, **, $P < 0.01$, vs SAL-sired male F1.

**Figure S3. Effects of METH$^{F1}$ administration on mPFC activity of naïve mice. a,** Experimental design and timeline. **b**, The mRNA levels of *c-Fos* and *Adrb1* in mPFC. **c**, The levels protein of c-Fos and ADRB1 in mPFC. S, saline-injected mice. M, METH-injected mice. N.S., $P > 0.05$ vs S mice.

**Figure S4. Effects of betaxolol on METH-preferred behavior in SAL-sired male F1. a,** Representative images of cannula location in mPFC. **b**, Timeline of Experimental S1. **c**, CPP score. **d**, ΔCPP score. **e,** Traveled traces in the CPP apparatus by SAL-sired male F1. **f**, Timeline of Experimental S2. **g**, CPP score. **h**, ΔCPP score. **i,** Traveled traces in the CPP apparatus by SAL-sired male F1. F1-SAL-Ts-Vcl, vehicle bilaterally into the mPFC of SAL-sired male F1 at 15 min prior to CPP test; F1-SAL-Ts-Bxol, betaxolol bilaterally into the mPFC of

SAL-sired male F1 at 15 min prior to CPP test; F1-SAL-Tr-Vcl, vehicle bilaterally into the mPFC of SAL-sired male F1 during CPP training; F1-SAL-Tr-Bxol, betaxolol bilaterally into the mPFC of SAL-sired male F1 during CPP training. N.S., $P > 0.05$. $^{##}$, $P < 0.01$ vs Pre-test CPP score. **, $P < 0.01$ vs F1-SAL-Ts-Vcl or F1-SAL-Tr-Vcl.

### Figure S1

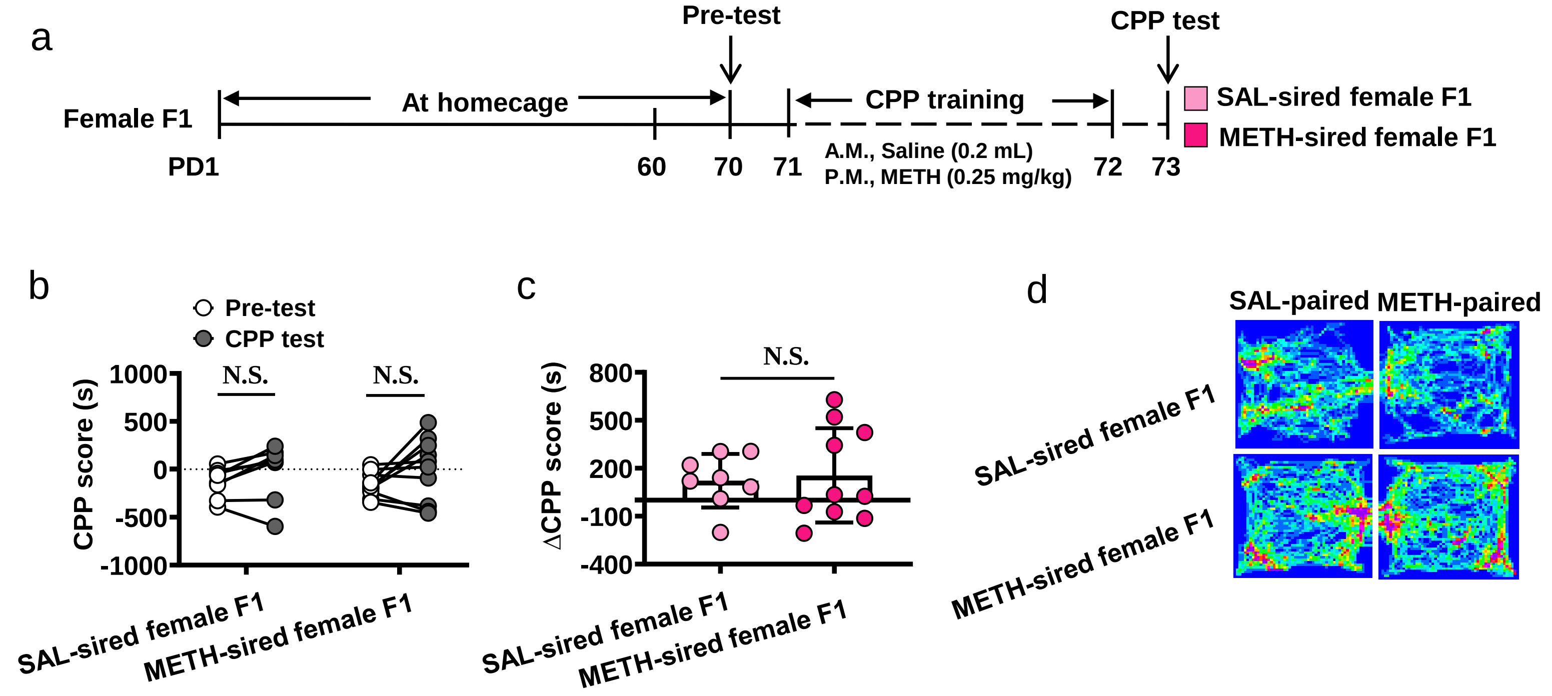

a
Pre-test
CPP test
Female F1
At homecage
CPP training
SAL-sired female F1
METH-sired female F1
PD1
60
70
71
A.M., Saline (0.2 mL)
P.M., METH (0.25 mg/kg)
72
73
b
Pre-test
CPP test
N.S.
N.S.
CPP score (s)
1000
500
0
-500
-1000
SAL-sired female F1
METH-sired female F1
c
ΔCPP score (s)
N.S.
800
500
200
-100
-400
SAL-sired female F1
METH-sired female F1
d
SAL-paired
METH-paired
SAL-sired female F1
METH-sired female F1

### Figure S2

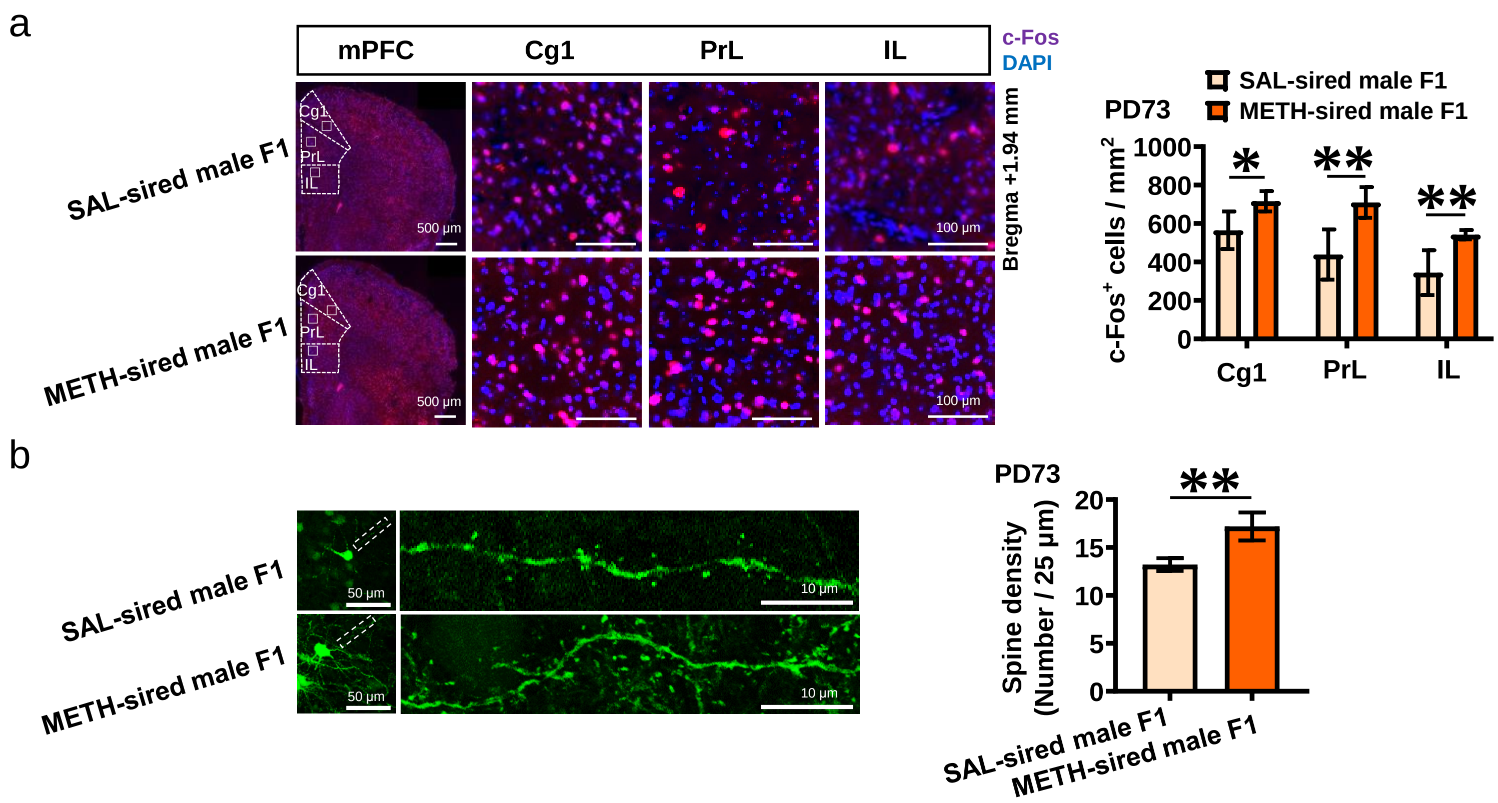
a
mPFC
Cg1
PrL
IL
c-Fos
DAPI
SAL-sired male F1
METH-sired male F1
Cg1
PrL
IL
500 μm
100 μm
Bregma +1.94 mm
PD73
SAL-sired male F1
METH-sired male F1
c-Fos+ cells / mm2
1000
800
600
400
200
0
Cg1
PrL
IL
b
50 μm
10 μm
PD73
Spine density
(Number / 25 μm)
20
15
10
5
0
SAL-sired male F1
METH-sired male F1

#### Figure S3

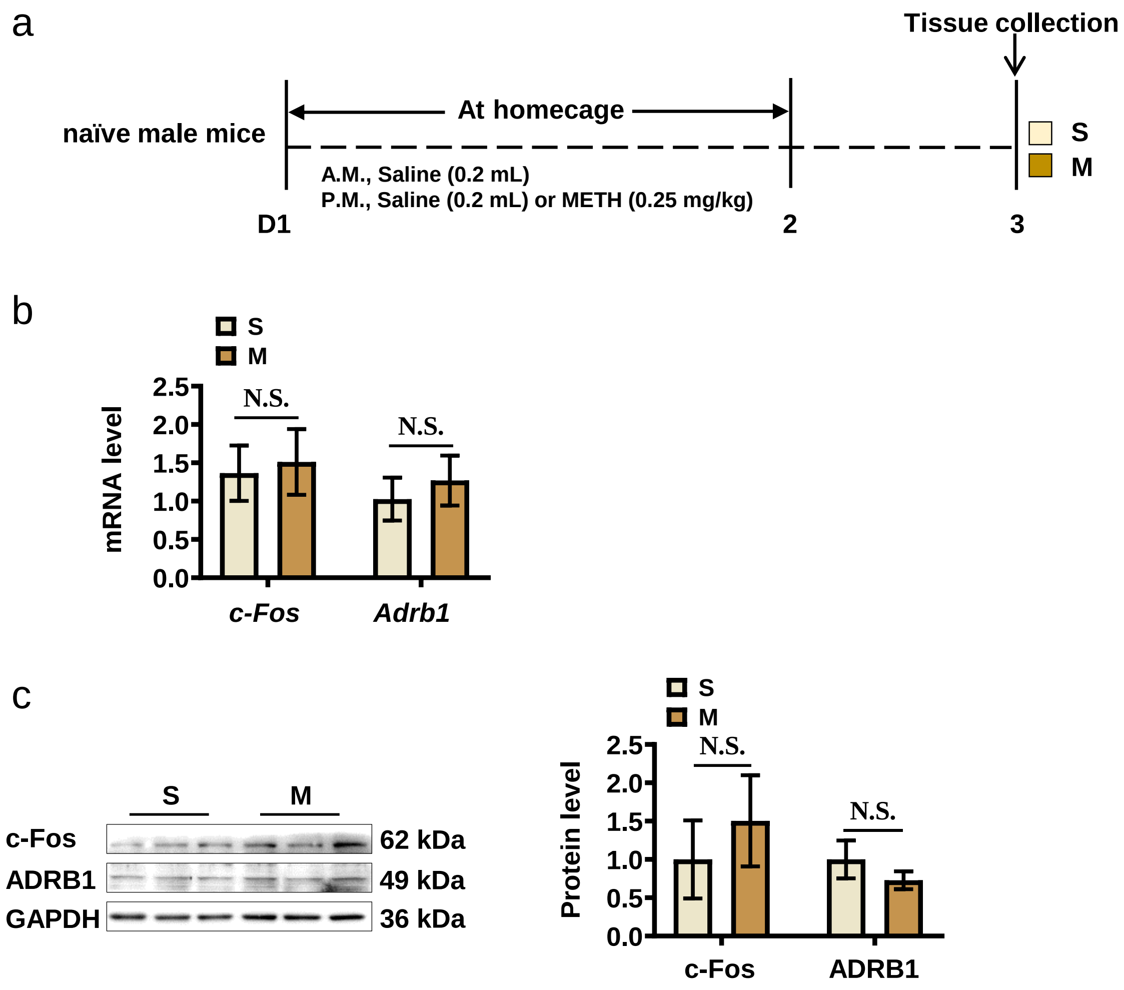

a
Tissue collection
naïve male mice
At homecage
A.M., Saline (0.2 mL)
P.M., Saline (0.2 mL) or METH (0.25 mg/kg)
D1
2
3
S
M
b
S
M
mRNA level
2.5
2.0
1.5
1.0
0.5
0.0
N.S.
N.S.
c-Fos
Adrb1
c
S
M
c-Fos
ADRB1
GAPDH
62 kDa
49 kDa
36 kDa
S
M
Protein level
2.5
2.0
1.5
1.0
0.5
0.0
N.S.
N.S.
c-Fos
ADRB1

### Figure S4

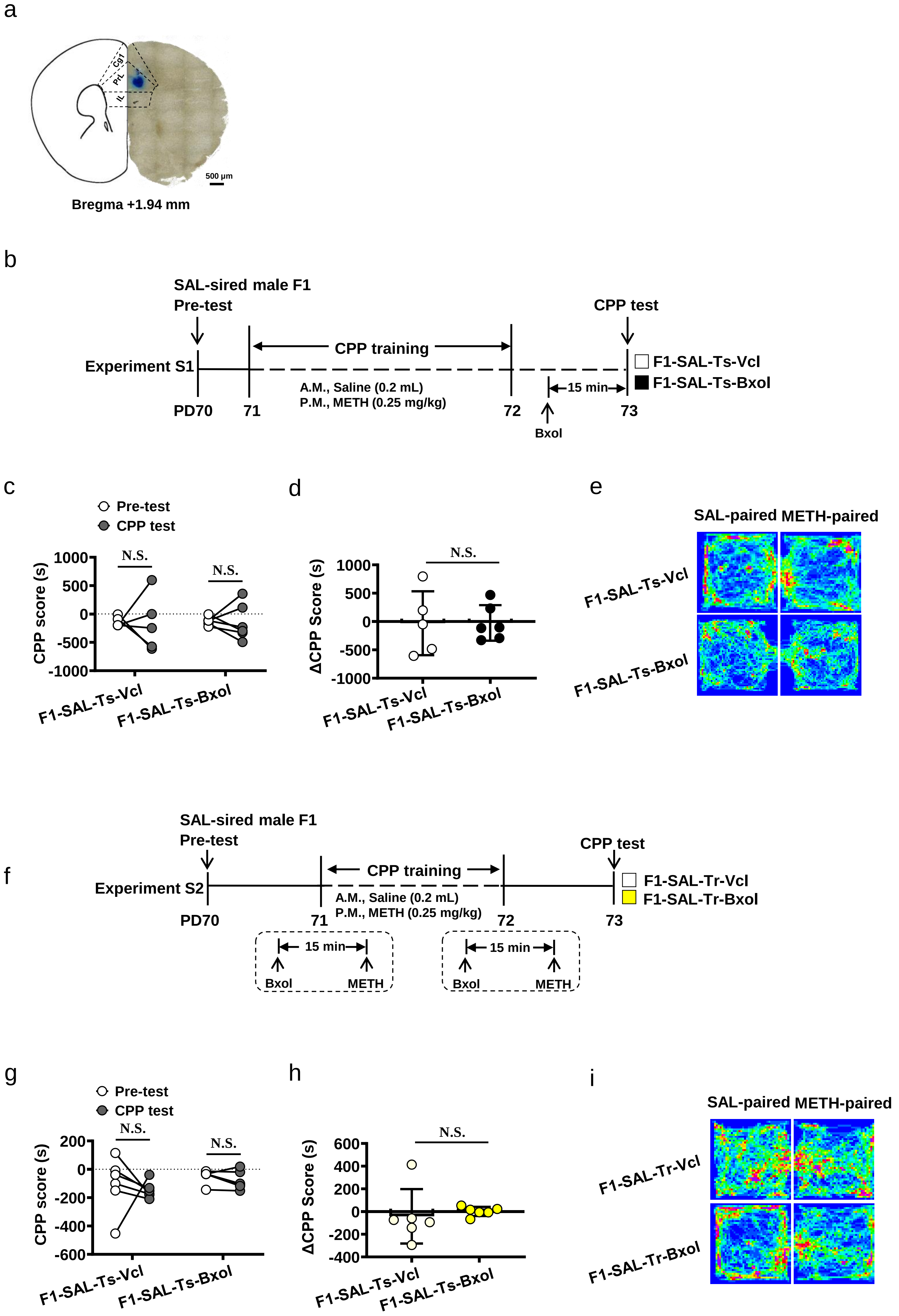
a
Cg1
PrL
IL
500 μm
Bregma +1.94 mm
b
SAL-sired male F1
Pre-test
CPP test
CPP training
Experiment S1
F1-SAL-Ts-Vcl
F1-SAL-Ts-Bxol
15 min
A.M., Saline (0.2 mL)
P.M., METH (0.25 mg/kg)
PD70
71
72
73
Bxol
c
Pre-test
CPP test
N.S.
N.S.
CPP score (s)
1000
500
0
-500
-1000
F1-SAL-Ts-Vcl
F1-SAL-Ts-Bxol
d
N.S.
ΔCPP Score (s)
1000
500
0
-500
-1000
F1-SAL-Ts-Vcl
F1-SAL-Ts-Bxol
e
SAL-paired
METH-paired
F1-SAL-Ts-Vcl
F1-SAL-Ts-Bxol
f
SAL-sired male F1
Pre-test
CPP test
Experiment S2
CPP training
F1-SAL-Tr-Vcl
F1-SAL-Tr-Bxol
A.M., Saline (0.2 mL)
P.M., METH (0.25 mg/kg)
PD70
71
72
73
15 min
Bxol
METH
15 min
Bxol
METH
g
Pre-test
CPP test
N.S.
N.S.
CPP score (s)
200
0
-200
-400
-600
F1-SAL-Ts-Vcl
F1-SAL-Ts-Bxol
h
N.S.
ΔCPP Score (s)
600
400
200
0
-200
-400
F1-SAL-Ts-Vcl
F1-SAL-Ts-Bxol
i
SAL-paired
METH-paired
F1-SAL-Tr-Vcl
F1-SAL-Tr-Bxol
